# Integrating transcription factor abundance with chromatin accessibility in human erythroid lineage commitment

**DOI:** 10.1101/2021.03.23.436609

**Authors:** Reema Baskar, Amy F. Chen, Patricia Favaro, Warren Reynolds, Fabian Mueller, Luciene Borges, Sizun Jiang, Hyun Shin Park, Eric T. Kool, William J. Greenleaf, Sean C. Bendall

**Affiliations:** Department of Pathology, Stanford University, Stanford, CA, USA; Cancer Biology Program, Stanford University, Stanford, CA, USA; Department of Genetics, Stanford University, Stanford, CA, USA; Department of Chemistry, Stanford University, Stanford, CA, USA; ChEM-H Institute, Stanford University, Stanford, CA, USA; Department of Applied Physics, Stanford University, Stanford, CA, USA; Chan Zuckerberg Biohub, San Francisco, CA, USA

**Author notes:** Contributed Equally (Reema Baskar, Amy F. Chen).

## Abstract

We present InTAC-seq, a method for simultaneous quantification of genome-wide open chromatin and intracellular protein abundance in fixed cells. Using InTAC we directly observe variation in chromatin accessibility and transcription factor motif occupancy driven by differences in transcription factor protein abundance. By purifying bone marrow progenitor cells based on GATA1 protein expression, we establish its role in both functional and epigenetic restriction of erythroid cell identity in human hematopoiesis.

## Main text

Transcription factors (TFs) are the master drivers of cell identity and differentiation, and their binding to specific regulatory sequences across the genome controls gene networks conferring cell phenotype and function. The identification of potential regulatory regions in a cell has been facilitated by techniques that map regions of open chromatin, such as the assay for transposase-accessible chromatin by sequencing (ATAC-seq)^1,2^. Still, methods to directly measure regulatory protein abundance in a cell and link changes in their levels to genome-wide changes in chromatin accessibility are lacking. Open questions in gene regulation remain around the relationship between TF abundance and its influence on accessibility and occupancy at predicted regulatory regions. The challenge of revealing these relationships is exacerbated when cells of interest are rare and regulatory proteins are transiently expressed in primary human tissue, such as hematopoietic lineage commitment in the bone marrow. However, by overcoming these challenges, measurements of protein abundance for lineage defining TFs in minimally manipulated primary tissues could be directly integrated with the anticipated restriction of epigenetic identity, providing unprecedented granularity into cell-fate decision mechanisms.

There have been a number of attempts to link gene expression to chromatin accessibility profiles employing single cell ATAC-seq (scATAC-seq) alone or in combination with RNA-seq in the same cell ^3-6^. While RNA levels are informative, they do not always reflect existing protein levels in the nucleus or account for post-translational regulation, which can dictate the functional state of a cell^7-9^. Protein expression has also been previously linked to chromatin accessibility in single cells (protein-indexed ATAC, or piATAC)^10^. However, TSS enrichment scores were considerably lower than those from conventional ATAC-seq libraries and piATAC datasets had comparatively few unique reads per cell. The lower enrichment in signal over background and decreased library complexity resulted in difficulty associating differences in TF protein abundance to significant changes in chromatin accessibility at predicted binding sites. Therefore, improvements in data quality will be critical in order to quantitatively assess how protein regulators (i.e. TFs, chromatin modifiers, upstream signaling molecules) influence epigenetic state.

Here, we introduce an improved method for intracellular staining and ATAC-seq (InTAC-seq, or InTAC) on fixed samples that produces libraries comparable in quality to those generated from fresh, unfixed cells. InTAC-seq can be applied to chromatin-binding protein factors such as TFs and generates high quality, quantitative data from primary tissue to robustly assess the relationship between TF abundance and chromatin occupancy. We first benchmarked InTAC-seq with the GM12878 lymphoblastoid cell line. We then further demonstrated that variation in GATA1 protein abundance differentially affects chromatin accessibility at sites of differing GATA1 binding affinity in the K562 erythroleukemic cell line. We then applied InTAC to bone marrow progenitor cells isolated based on GATA1 expression to profile the GATA1-associated epigenetic changes in human hematopoietic progenitor differentiation. The high-quality profiles produced by InTAC enabled us to integrate our results with previous single cell ATAC- and RNA-seq bone marrow datasets. Thus, we could position the isolated GATA1 cells within this multiomic landscape of human hematopoietic lineage commitment. Our results reveal that while GATA1 is expressed in only a subset of conventional erythroid progenitors (i.e. the megakaryocytic and erythroid progenitor population, or MEPs), it almost entirely captures the erythropoietic program. By using surrogate surface markers to enrich for GATA1 expression we show that GATA1-high progenitor cells demonstrate the most functionally pure erythroid progenitor capacity identified to date. Thus, our data define GATA1 as a central epigenetic and functional lineage restriction factor in human red blood cell homeostasis.

The main challenge with combining ATAC-seq with intracellular protein quantification lies in the fixation and permeabilization of cells required for direct measurement of intracellular proteins using affinity reagents such as antibodies and subsequent paraformaldehyde crosslinking reversal at high temperature prior to library amplification. The conventional 65°C crosslink reversal step^10^ results in dissociation and loss of shorter fragments that contribute to a large fraction of the final ATAC-seq library, resulting in reduced library complexity and lower TSS enrichment scores. To improve data quality, the InTAC protocol uses a shorter fixation time with mild permeabilization and reversal of formaldehyde crosslinks with a catalyst^11^ to allow crosslink reversal to occur at 37°C (Figure 1A).

**Figure 1.**
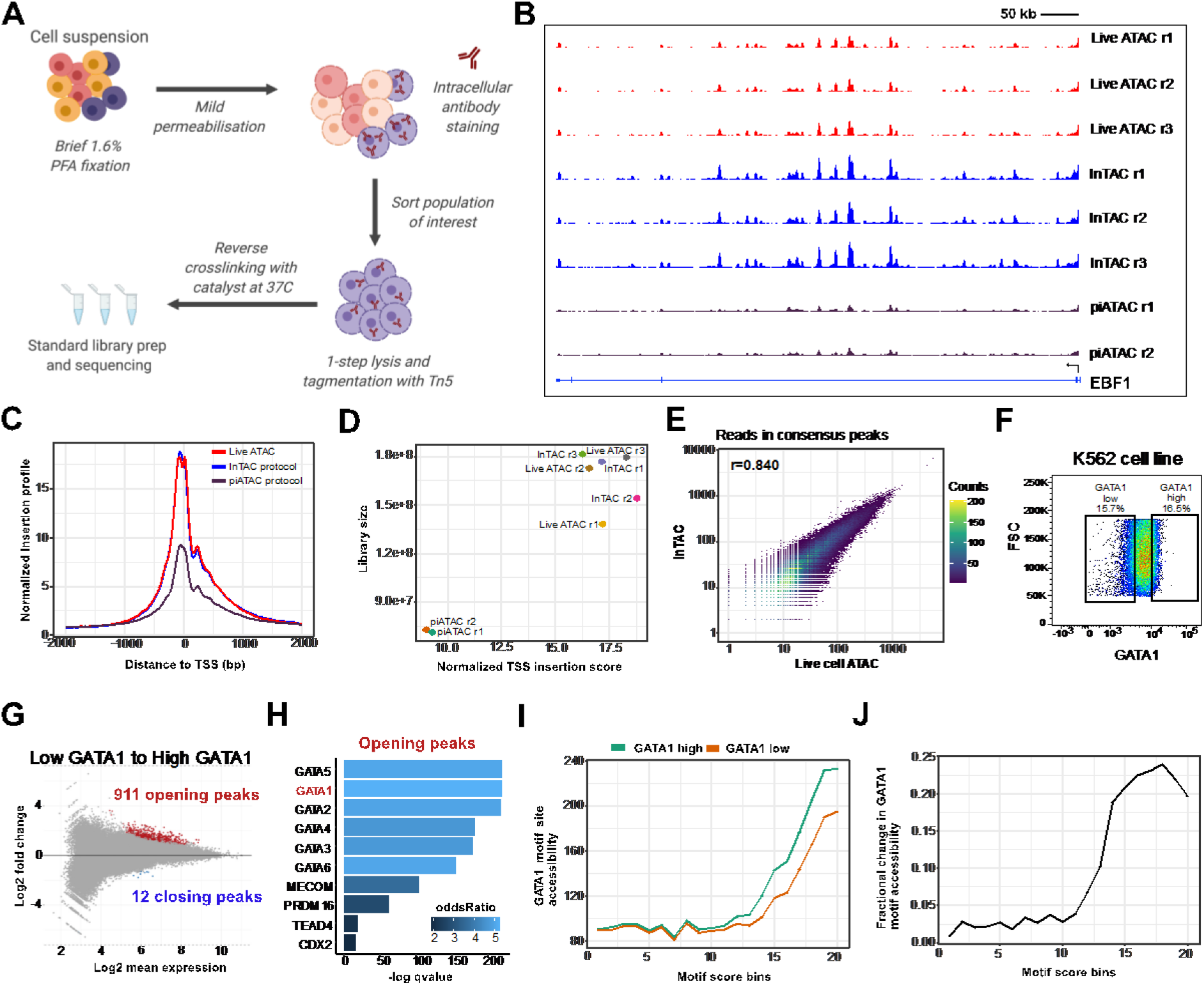
InTAC data from fixed cells is of comparable quality to ATAC-seq data from live cells and allows interrogation of TF chromatin occupancy as a function of its protein abundance **A**. Overview of InTAC experimental protocol. **B**. Genome coverage of ATAC-seq data generated from live cells, fixed cells using InTAC, or fixed cells using piATAC at the EBF1 locus in GM12878 cells. **C**. Normalized Tn5 insertion profiles centered at transcription start sites (TSS) for the indicated ATAC-seq libraries. **D**. Scatterplot of estimated library size vs normalized TSS insertion score across all replicates of compared protocols. **E**. Scatterplot of reads in consensus peaks averaged across replicates between InTAC and live ATAC samples, with calculated Spearman correlation coefficient as shown. **F**. FACS plot of forward scatter (linear scale) vs GATA1 protein abundance (log10 scale) and the gating strategy to isolate the highest and lowest 15% of GATA1-expressing K562 cells. **G**. MA plot of log2 fold change in accessibility between GATA1-high and GATA1-low K562 populations vs log2 mean number of reads at all consensus peaks. Peaks with significant changes in accessibility are highlighted in red or blue. **H**. Most significantly enriched TF motifs in differentially accessible peaks in GATA1-high cells calculated using Fisher’s test. **I**. Average accessibility of GATA1 motif sites across all consensus ATAC-seq peaks binned by GATA1 motif score. Accessibility is defined here as the area under the curve of a plot of bias-corrected, normalized Tn5 insertions centered at GATA1 motif sites (as in Fig S1G), integrated from −50 to +50 bp and excluding the TF footprint from −10 to +10 bp. **J**. Difference in GATA1 motif accessibility between GATA1-high and GATA1-low samples normalized to the accessibility in the GATA1-low population for each motif score bin.

ATAC-seq libraries generated from fixed GM12878 cells using InTAC exhibited enrichment in Tn5 insertions at transcriptional start sites (TSS) comparable to the TSS enrichment in ATAC-seq libraries from unfixed live cells and approximately two-fold greater than that in published bulk piATAC libraries (Figure 1C). The fragment length distribution for InTAC libraries was also similar to the distribution in libraries from live cells due to preservation of sub-nucleosomal fragments at 37°C (Figure S1A). The retention of sub-nucleosomal fragments translated into a larger estimated number of unique ATAC-seq fragments from InTAC and live ATAC libraries relative to piATAC libraries (Figure 1D). To compare InTAC libraries to unfixed libraries, we plotted the correlation of reads in consensus peaks and found a strong correlation between InTAC and live samples (Figure 1E, S1B). Genome browser tracks further illustrated the concordance between InTAC and live samples (Figure 1B). We further demonstrated that InTAC can be performed on as low as 100 cells, enabling profiling of rare cell types (Figure S1C and D). Together, these results show that the InTAC protocol produces ATAC-seq libraries that closely resemble live ATAC-seq libraries, with comparable signal-to-background (as defined by TSS enrichment scores) and library complexity.

We next aimed to use InTAC to detect chromatin accessibility differences in cells endogenously expressing different levels of a chromatin-binding protein. We compared the chromatin accessibility profiles in K562 cells expressing the highest or lowest 15% of GATA1 levels using InTAC (Figure 1F). With ChromVAR^12^, we found that GATA binding motifs were among the most variable in accessibility across samples (Figure S1E). Specifically, we observed increased accessibility in cells with high levels of GATA1 at GATA1 motif sites (Figure S1G and H). Differentially accessible peaks (FDR < 0.1) between cells with high vs low GATA1 were almost exclusively more accessible in GATA1-high cells, and these were most significantly enriched for GATA motifs relative to all other peaks present across samples (Figure 1H, S1F), suggesting that accessibility differences are due to differences in GATA1 abundance.

We next asked how increases in GATA1 occupancy can vary based on the predicted binding affinity of the GATA1 binding site. To address this, we grouped all consensus peaks into 20 bins of equal size based on the quality of their GATA1 motif score and measured the average accessibility across these bins in cells with high vs low GATA1. We found that cells with high GATA1 levels exhibit a general increase in accessibility at GATA1 motif sites above a threshold motif score that likely represents the minimum score for a true GATA1 binding site (Figure 1I). However, we observed that the greatest change in accessibility between cells with high vs low GATA1 occurred at sites of moderate predicted affinity (i.e. bin 18), suggesting that the occupancy of GATA1 at the highest affinity GATA1 binding sites may be saturated even at low GATA1 levels (Figure 1J). These results showcase the ability of InTAC to measure subtle differences in chromatin accessibility among populations, allowing us to link natural variation in transcription factor abundance to functional differences at the chromatin level and approximate TF occupation stoichiometry.

While clinically significant for both cell-based therapies and hematopoietic dysplasias, the regulatory landscape of erythropoietic cell homeostasis in the human blood stem cell compartment is not well understood. Bone marrow progenitor populations are traditionally defined and isolated based on expression of cell-surface proteins, which viably preserves these cells for downstream functional assays^13^. However, these surface molecules are not always functionally related to the cellular state that we associate them with. We therefore hypothesized that intracellular regulatory protein abundance would identify cellular states with higher fidelity and enable more accurate molecular characterization. To test this, we focused on human erythroid progenitor cells, which are regulated by GATA1^14^ but conventionally defined by unrelated surface molecules (i.e. IL3 receptor CD123, and pan-leukocyte phosphatase isoform CD45RA) to delineate megakaryocyte erythroid progenitor (MEP) cells. We applied InTAC to interrogate the link between the abundance of the lineage-defining TF, GATA1, and epigenetic commitment to red blood cell development.

First, we used InTAC to profile the accessible chromatin landscape of GATA1-positive and GATA1-negative cells within the landscape of the general bone marrow progenitor compartment (i.e. CD34^+^CD38^+^) to determine if GATA1-positive cells were enriched for erythroid epigenetic signatures (Figure 2A). InTAC libraries generated from these isolated subpopulations had TSS enrichment scores similar to ATAC-seq libraries from live GM12878 cells indicating that InTAC performs well on primary human samples (Fig S2A and B). We observed a marked increase in accessibility at regulatory regions surrounding the GATA1 locus in GATA1-positive vs GATA1-negative progenitors consistent with GATA1 expression levels, along with a decrease in accessibility at regulatory regions within the SPI1 locus, which encodes a TF known to antagonize GATA1 activity and repress MEP commitment^15-18^ (Figure 2B). The binding motifs for these two TFs also exhibited strong differences in accessibility between GATA1-positive and negative progenitors (Figure 2C). We further observed a broader trend of increased accessibility surrounding the motifs for TFs that drive erythroid fates (eg. GATA1, Mecom^19^) and decreased accessibility at motifs for TFs that drive myeloid and lymphoid lineages (eg. SPI1^20,21^, EBF1^22^) in GATA1-positive progenitors (Figure 2C). Analysis of differentially accessible peaks showed enrichment of these erythroid TF motifs in sites more accessible in GATA1-positive progenitors and enrichment of a variety of myeloid and lymphoid TFs in sites more accessible in GATA1-negative progenitors (Figure 2D and E, S2C). Deeper analysis into accessibility at GATA1 motif sites of varying binding affinity across the genome showed that high GATA1 abundance in progenitors is associated with marked increase in accessibility above a threshold motif score (Figure S2D). The greatest increases were observed at sites with the highest predicted GATA1 affinity, confirming the preferential binding of GATA1 to the genome at these sites (Figure S2E). Altogether, these data suggest that GATA1 expression in human hematopoietic progenitors promotes an erythroid epigenetic program while repressing alternative fates.

**Figure 2.**
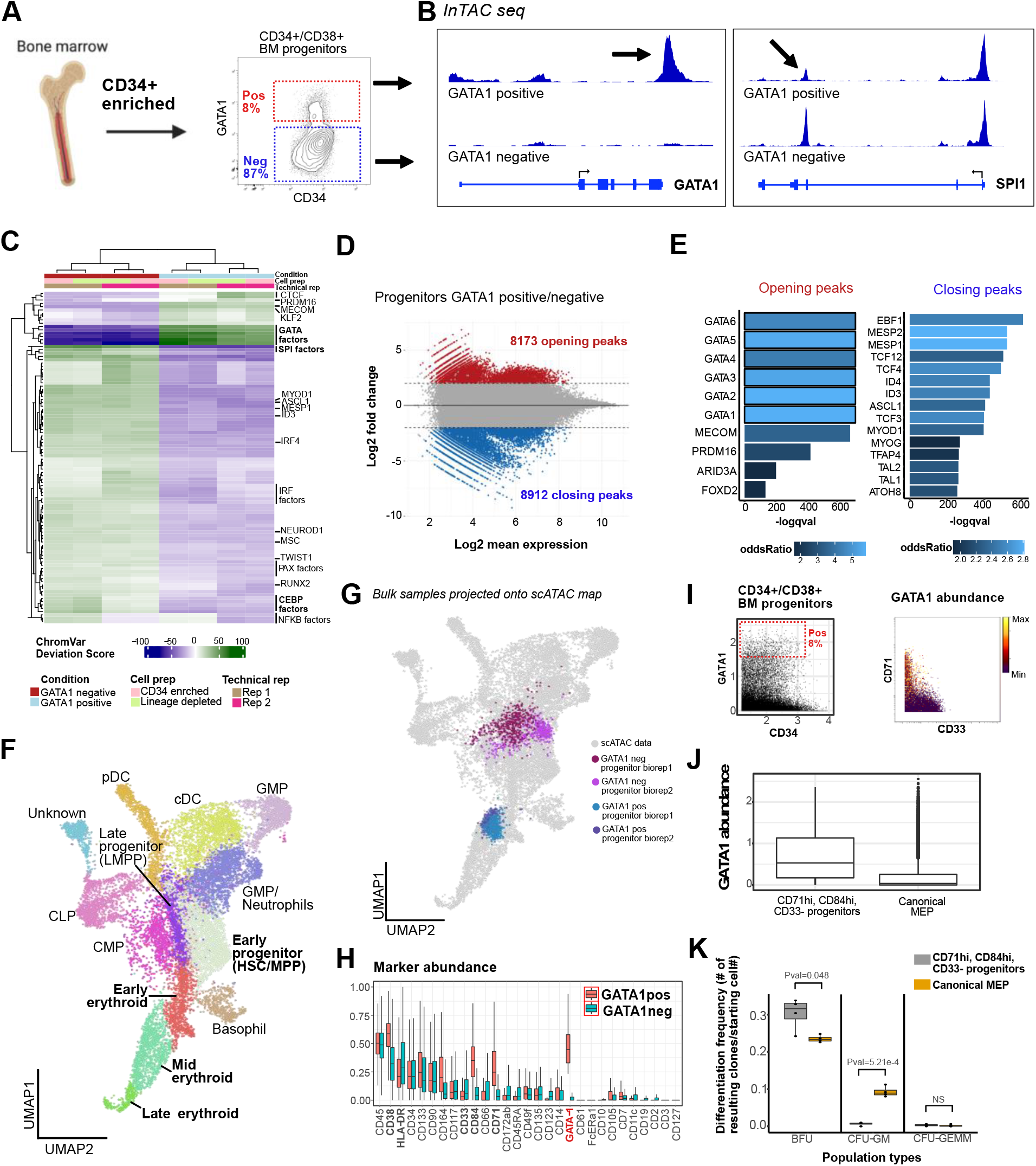
GATA1-high BM progenitors are enriched for erythroid potential **A**. Bone marrow aspirate is ficolled and enriched for CD34+ cells before gating for CD34+/CD38+ cells and selecting GATA1 positive population (top ∼8%) and negative population (bottom ∼87%). **B**. InTAC genome coverage plots at GATA1 and SPI1 loci for GATA1 positive and negative BM progenitors. **C**. Heatmap of chromVAR deviation scores across GATA1 positive and negative BM progenitors for top 50 most variable motifs. **D**. MA plot of log2 fold change in accessibility between GATA1 positive and negative BM progenitors vs log2 mean number of reads in consensus peaks. Peaks with significant changes in accessibility are highlighted in red or blue. **E**. Most significantly enriched TF motifs in differentially accessible peaks between GATA1 positive and negative BM progenitors calculated using Fisher’s test. **F**. UMAP of previously published and annotated BM scATAC datasets (Grenja et. al.) with Seurat clusters manually annotated as key BM populations. **G**. Bulk BM progenitor InTAC data simulated as single cell ATAC counts and projected onto scATAC UMAP space. **H**. Normalized surface marker abundance of GATA1 positive BM progenitors (top ∼8% of expression) and GATA1 negative BM progenitors (bottom ∼87% of expression) from mass cytometry **I**. (left) Scatter plot of GATA1 by CD34 protein expression where GATA1 positivity in cytometry space is defined as top 8% of expressing cells. (right) Scatter plot of GATA1-enriching marker CD71 vs depleting marker CD33 colored by GATA1 abundance in BM progenitor space. **J**. GATA1 protein abundance in manually gated target population (as defined by CD71hi, CD84hi, CD33neg) versus canonically gated MEP population. **K**. Clonal differentiation frequency of target population and canonical MEP population to different lineages/population types.

To further assess the position of these progenitors within the hematopoietic hierarchy, we projected our GATA1 progenitor samples onto a single cell ATAC-seq UMAP of filtered bone marrow mononuclear cells (BMMCs) from Granja et al.^4^ to identify their closest hematopoietic cell type using ArchR^23^. Given our focus on erythropoiesis, we performed a more fine-grained analysis of the erythroid cluster and separated it into early, mid, and late erythroid progenitor populations based on differences in gene accessibility and chromVAR deviation scores of key TFs (Figure 2F, S2F, G, and H). We found that GATA1-positive progenitors were positioned within the mid erythroid progenitor branch while GATA1-negative progenitors spanned annotated GMP, LMPP, and CMP clusters (Figure 2G). This observation is consistent with the expected presence of GMPs, LMPPs, and CMPs in CD34+CD38+ bone marrow cells^13^. More strikingly, the restriction of GATA1-positive progenitors to the mid erythroid cluster (Figure 2F and G) combined with the overall enrichment of erythropoietic programs (Figure 2C and D) suggests that progenitor expression of GATA1 alone is likely sufficient to enrich for erythroid commitment.

To functionally assess erythroid progenitor commitment in GATA1-expressing cells, we sought to benchmark them against conventional human MEPs (CD34+CD38+CD10-CD123-CD45RA-) using a colony-forming assay for clonal hematopoietic lineage potential. In order to isolate viable cells enriched in GATA1 for the assay, we first identified surface protein surrogates for GATA1 expression. To identify GATA1 cell surface correlates, we co-stained BMMCs with metal isotope-conjugated antibodies to GATA1 and a 30-plex panel of relevant surface proteins in BM for interrogation using mass cytometry^24^ (Figure S2I and Table 1). We found that CD71 and CD84, both known to be enriched in erythroid cells^25^, best represented GATA1 expression (Pearson correlation = 0.372 and 0.378, respectively; Figure S2J). The myeloid-enriched surface protein, CD33, was also mildly anti-correlated with GATA1 (Pearson correlation = −0.079, Figure S2J). Additionally, when we selected the high expressing GATA1-positive progenitors (Figure 2I left) at a frequency similar to that used to sort for GATA1-positive progenitors in the InTAC experiment (∼8%), we observed an enrichment for CD71, CD84, and CD33 in the GATA1-positive relative to GATA1-negative populations (Figure 2H). Lastly a data-driven, back-gating approach was employed^26^ and CD33, CD84 and CD71 were predicted to best enrich for the GATA1 positive target population with high F-score at 0.99 (red cells, Figure S2K, L). By selecting for CD71+CD84+CD33-cells within the CD34+CD38+ compartment, we confirmed that these cells have higher expression of GATA1 relative to cells isolated with conventional MEP markers (Figure 2J).

We then compared the hematopoietic colony-forming potential of the CD71+CD84+CD33-(i.e. GATA1-enriched) population and conventional MEPs using a methylcellulose assay. The CD71+CD84+CD33-(GATA1-enriched) cells were significantly (p=0.048) enriched for erythroid (BFU-E) colonies compared to conventional MEPs that formed a significant number of granulocyte and monocyte (CFU-GM) colonies (p=5.21E-4, Figure 2K and S2M), which is consistent with previous reports^27,28^. These data confirm that GATA1 expression in human CD34+CD38+ progenitors is sufficient to identify erythroid commitment, and is consistent with the GATA1 epigenetic programs we observed.

Consistent with their mixed lineage potential demonstrated by CFU analysis, conventional human MEPs have mixed positive expression for GATA1 (Figure 3A). To examine the nature of GATA1-high cells from within the conventional MEP population and how they compare to the GATA1-positive CD34+CD38+ progenitors, we isolated them for InTAC (Figure 3A, S3A). Despite the low cell numbers obtained from these rare primary BM populations (down to ∼250 cells), the resulting ATAC-seq data were of high quality (Figure S3B). As expected, projection of GATA1-high cells isolated both from CD34+CD38+ progenitor and MEP compartments onto the scATAC UMAP of BMMCs demonstrated similar embedding within the mid-erythroid population (Figure 3B). These data suggest that they are, in fact, similar cells and further confirming that GATA1 expression in BM progenitors is sufficient to isolate erythroid progenitors, including as a subset of the conventional MEP population (Figure 3B).

**Figure 3.**
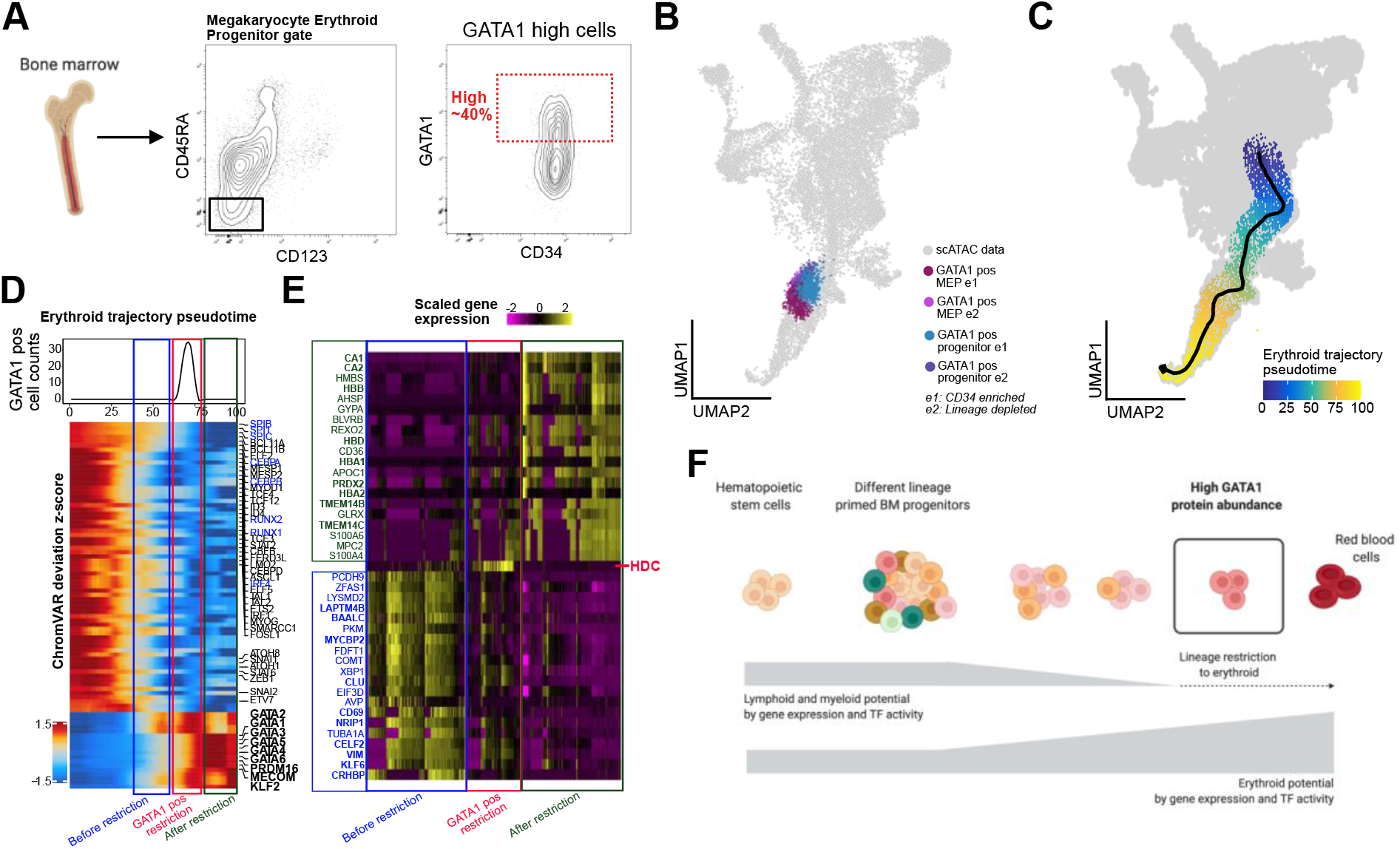
GATA1 protein abundance delineates key erythroid commitment step in RBC development **A**. Bone marrow aspirate is ficolled and enriched for CD34+ cells before gating for conventional MEP (CD34+/CD38+/CD10-/CD45RA-/CD123-) cells and selecting highest (25-40%) GATA1-expressing MEP and remaining low GATA1-expressing MEP cells. **B**. Bulk GATA1 positive BM progenitor and MEP InTAC data simulated as single cell ATAC counts and projected on scATAC UMAP space. **C**. Putative erythropoiesis trajectory constructed from HSC to late erythoid population and overlaid on scATAC UMAP. **D**. Heatmap of top variable TFs by ChromVAR deviation scores across constructed erythroid trajectory with the projected position of GATA1-positive samples indicated in red as GATA1 pos restriction, in blue as before restriction and in green as after restriction. Top: Line plot of GATA1 pos scATAC cells as binned across pseudotime **E**. Top 20 genes significantly enriched (of fold change 2 and above) in integrated scRNA seq data between the 3 bins, before, during and after GATA1 positive restriction. **F**. Summary schematic of continuous differentiation to erythrocytes in BM with lineage restriction from downregulation of lymphoid/myeloid TF activity and gene expression programs and upregulation of erythroid TF activity and gene expression programs. GATA1 protein abundance marks key restriction point of erythroid lineage commitment.

During erythropoiesis, BM progenitors undergo global changes in transcription factor activity and chromatin accessibility in order to restrict non-erythroid lineage potential and drive the erythroid program. In order to model this process within our data and understand the epigenetic transitions associated with GATA1 acquisition, and thus erythroid commitment, we calculated a pseudotime ordering of cells from HSCs to the late erythroid cluster (Figure 3C). Using the GATA1 cells as a landmark, we identified the positions of their closest scATAC BM cells within the erythroid trajectory (Figure S3H). To assess the chromatin accessibility changes across erythroid differentiation, we then plotted chromVAR deviations for variable TF motifs along this derived erythroid trajectory. We observed a coordinated decrease in accessibility at motif sites for TFs that drive alternative lineages and an increase in accessibility at erythroid TF motif sites precisely where our GATA1-positive progenitors are positioned within the trajectory (Figure 3D, red box and erythroid TFs in bold). The preceding stage of erythroid differentiation still shows persistence of non-erythroid TF motif accessibility (Figure 3D, blue box), consistent with functional lineage restriction and erythroid fate determination being tied to GATA1 protein abundance.

Using existing scRNA-seq data from the same BM samples, we next integrated measured TF protein abundance data with gene expression, gene accessibility (gene score), and motif accessibility along a differentiation trajectory. We observe that gene accessibility changes before and after our determined GATA1-positive erythroid restriction point concur with expected trends for lymphoid/myeloid and erythroid genes, respectively (Figure S3J). Differential analysis of integrated scRNA-seq data at these 3 stages of erythroid differentiation reveal key hemoglobin subunit genes (eg. HBD, HBA1, HBB) and heme metabolism (TMEM14B/C) after the restriction point (Figure 3E, in green and red). We also observe differential upregulation of histidine decarboxylase (HDC), an enzyme in the histamine synthesis pathways, in cells at this restriction point, with decreasing HDC expression as cells move into the final erythroid stage. In contrast, preceding the GATA1-defined restriction point, we see enrichment for genes involved in various non-erythroid lineages (Figure 3E, in blue, with genes related to hematopoiesis bolded), including NRIP1 for hematopoietic stem cell quiescence^29,30^. Integrated maps further reveal the discord between the different regulatory layers for specific genes and highlight the importance of directly measuring protein levels of key TFs and associated chromatin accessibility due to the presence of TF families that recognize similar binding motifs (Figure S3I).

Given that conventional MEPs have mixed GATA1 expression (Figure 3A) and corresponds to mixed commitment to the erythroid lineage (Figure 2K), we also examined how the GATA1-high and GATA1-low MEP cells differed. We observed differences in accessibility at the gene locus for the erythropoiesis-enhancing TF, Myb, a known target of GATA1 binding and repression, between these two populations, consistent with their GATA1 levels (Figure S3C). Notably, we found that GATA1 expression marked distinct cellular subsets with thousands of differentially accessible sites between GATA1-high and GATA1-low MEPs (Figure S3D, E). While megakaryocyte and erythroid TF motifs were enriched in GATA1-high cells, motifs for TFs involved in myeloid and B cell development were enriched in GATA1-low cells (Figure S3F, G). These results suggest that GATA1-low cells within the conventional MEP population retain some myeloid and/or lymphoid potential, likely resulting in their mixed lineage functional output in clonal assays. Interestingly, these observations are consistent with a previous study that described a subpopulation of cells within the MEP compartment exhibiting both erythroid and myeloid potential based on functional assays, although lymphoid potential was not tested^31^.

In summary, we have developed a robust protocol for profiling chromatin accessibility in fixed cells that enables staining and isolation of populations based on protein levels of intracellular regulators prior to ATAC-seq. This protocol, which we termed InTAC-seq, enables us to integrate endogenous differences in key transcription factors with associated chromatin accessibility profiles to directly probe the link between their protein level abundance and chromatin occupancy. We use our approach to reveal GATA1-associated epigenetic profiles and infer motif binding stoichiometry.

Importantly, our protocol captures high quality data from primary human tissue and can be used with low (∼100 cells) inputs which allows direct and robust integrated profiling of intracellular functional drivers and their associated chromatin accessibility in rare populations such as progenitors in human bone marrow. We revealed the GATA1-associated epigenome in CD34^+^CD38^+^ hematopoietic progenitors and identified strong epigenetic restriction to erythroid cell fate. We further demonstrated that, as predicted based on their epigenetic restriction, these GATA1-expressing cells represent the best example of erythroid functional restriction compared to previous definitions. Consequently, GATA1 expression differences within MEP likely account for their lineage heterogeneity, given residual myeloid and/or lymphoid potential in GATA1 low MEPs.

InTAC enables biologists to directly connect levels of functionally pertinent intracellular proteins such as transcription factors and other chromatin-binding proteins to chromatin accessibility profiles in order to answer gene regulatory questions and assess transient cellular molecular states previously difficult to access. By measuring endogenous intracellular regulators in complex biological contexts, InTAC delineates epigenetic landscapes driven by master TFs or chromatin remodelers in primary human tissue. This allows us to interrogate how levels of these key proteins and their cooperative and antagonistic partners could influence global chromatin accessibility and local binding to drive cellular state and function. Additionally, one could devise more complex isolation schemes based on the expression of a multitude of intracellular regulators, rather than those which could be accomplished with more targeted chromatin immunoprecipitation (ChIP) procedures if sufficient cell numbers were available. Such an approach could identify cooperative or antagonistic functions with the key protein(s) of interest and associate their expression levels with specific cell states. Alternatively, we can design experiments that introduce exogenous factors into biological systems and use InTAC to isolate and profile populations with a range of expression of the exogenous proteins to understand the relationship between their abundance and chromatin architecture. InTAC is also complementary to single cell genomic techniques such as single cell ATAC-seq and RNA-seq as it can be integrated together with existing single cell data so that the populations of interest can be studied in the context of a larger biological system. Overall, InTAC enables the profiling of chromatin accessibility of cell populations isolated by abundance of intracellular proteins such as transcription factors with a cost, ease, and robustness that is on par with standard ATAC-seq.

## Methods

### Cell lines

The human chronic myeloid leukemia cell line, K562, were obtained from the American Type Culture Collection (Manassas, VA, USA). The human lymphoblastic cell line GM12878 were obtained from Coriell Institute. Cells were cultured in RPMI 1640 medium containing 15% fetal bovine serum (FBS) and penicillin/streptomycin, and maintained at 37 °C, 5% CO_2_.

### Primary bone marrow samples

All fresh adult whole BM used in this study was collected in heparin sulfate anticoagulant and purchased from All Cells, Inc. BM mononuclear cells (BMMNC) were separated using Ficoll-Paque Plus (Amersham Biosciences). Next, BMMNC were either cryopreserved in FBS with 10% of DMSO or previously enriched for CD34^+^ (CD34 MicroBead Kit, Miltenyi Biotec) before cryopreservation.

### K562 GATA-1 staining and sorting

Cells were washed once with PBS, then incubated in LIVE/DEAD™ Fixable Aqua Dead Cell Stain (Invitrogen L34965) diluted in PBS for 30 mins on ice. After a PBS wash, cells were fixed with 1.6% formaldehyde in PBS for 1 min, then quenched with an equal volume of 1X eBioscience permeabilization buffer (Thermofisher Scientific 00-8333-56). Cells were immediately centrifuged for 5 min at 600g and washed once with permeabilization buffer. Cells were then stained with anti-cleaved caspase 3-PE (BD #550821) and anti-GATA1 (Abcam ab181544) for 30 mins at room temperature and FITC anti-rabbit secondary (Cell Signaling Technologies #4412S) for 30 mins. Washes were performed using permeabilization buffer and cells were resuspended in PBS for FACS and sorted using a BD FACSAria II. Cells positive for Aqua live/dead stain or cleaved caspase 3 were gated out and a narrow FSC gate was used to control for cell size. Cells in the lowest and highest 15% of GATA1 expression were then sorted into PBS containing 30% FBS for InTAC.

### Bone marrow processing and GATA-1 sorting

On the day of the sorting, BM enriched CD34^+^ cells or BMMNC were thawed into cell culture medium supplemented with 25 U/mL benzonase (Sigma-Aldrich). For BMMNC, cells underwent magnetic lineage depletion according to the manufacturer’s instructions using BD Streptavidin Particles Plus (BD Biosciences #557812) and the BD IMag Cell Separation Magnet (BD Biosciences) with biotinylated anti-CD3, CD15, CD7, and CD56. Next, cells were incubated with LIVE/DEAD fixable Aqua dead cell stain (Invitrogen #L34957) for 30 minutes at room temperature (RTP) in dark, followed by a wash with PBS, before Fc receptor blocking (Human TruStain FcX, Biolegend #422302) for 10 minutes. Surface staining was carried out with CD34-FITC, CD38-BV421, CD45RA-AlexaFluor700, CD10-BV650, and CD123-PECy7 at 4°C in dark. Cells were then washed with cell staining media (PBS + 0.5% BSA). Fixation was carried out with 1.6% paraformaldehyde for 5 min at RTP before quenching with 1X eBioscience permeabilization buffer (Thermo Fisher Scientific 00-8333-56). Two washes were carried out with permeabilization buffer at 600G for 5 minutes each. Cells were incubated with GATA1-PE in permeabilization buffer for 45 minutes at RTP, followed by a wash with CSM and sorting using a BD FACS Aria II (BD Biosciences). GATA-1 high and low/intermediate gates were performed on BM progenitors (gated as singlet, viable CD34^+^, CD38^+^ cells) and on conventional MEP (gated as singlet, viable, CD34^+^, CD38^+^, CD10^−^, CD123^−^, CD45RA^−^ cells; antibody panels in Supplementary Table 2).

### InTAC protocol

Fixed, permeabilized cells were counted using a hemocytometer and up to 50,000 cells were used for ATAC-seq where possible. Cells were spun down at 600g for 5 mins and resuspended in transposition mix containing 1X TD buffer, 0.1% NP40, 0.01% digitonin, and Tn5. Cells were incubated at 37 degrees with 1200 rpm shaking for 1 hour. 2X reverse crosslinking buffer (2% SDS, 0.2mg/mL proteinase K, and 100mM N,N-Dimethylethylenediamine, pH 6.5 [Sigma Aldrich D158003]) was added at equal volume to transposed cells and reversal of crosslinks was performed at 37 degrees overnight with 600 rpm shaking. DNA was purified using Qiagen minelute PCR purification columns and ATAC-seq libraries were generated as previously described in Buenrostro et al. 2015. For preparation of live ATAC-seq samples from fresh cells as a comparison, samples were prepared as above, except DNA was purified immediately following transposition rather than performing crosslink reversal.

### ATAC-seq data processing

Adapters were trimmed using cutadapt and reads were mapped using bowtie2 with max fragment length of 2000bp to hg19 (primary bone marrow samples) or hg38 (all cell lines). We then filtered for non-mitochondrial reads, mapq > 20, and properly paired reads. We then removed duplicates using Picard tools. Peaks were called using macs2 with the following parameters on Tn5 insertion sites: ╌shift −75 ╌extsize 150 ╌nomodel ╌call-summits ╌nolambda -p 0.01 -B – SPMR

### ATAC-seq QC of live and fixed GM12878 samples

For estimating library complexity, libraries were downsampled to 13 million read pairs prior to deduplication and library size was estimated using Picard tools EstimateLibraryComplexity. For TSS enrichment, deduplicated libraries were down-sampled to 10 million read pairs except for cell titration experiments where libraries were down-sampled to 4 million read pairs. TSS enrichment was calculated using the getTssEnrichment function in the ChrAccR R package for gencode.v27 protein coding gene transcriptional start sites.

### Differential accessibility analysis

Aligned, deduplicated bam files output from data processing pipeline were loaded into R in HDF5 format using DsATAC.bam function in the ChrAccR R package. Consensus peakset across technical and biological replicates was calculated using getPeakSet.snakeATAC function in the ChrAccR R package where peaks have to be consistently absent or present across replicates to be retained. Count matrix was calculated as insertion counts across samples at consensus peakset regions using ChrAccR regionAggregation function. DESeq2^32^ was used to calculate differentially accessible peaks and independent hypothesis weighting (cite IHW) was used to correct for multiple testing. ggmaplot package was used to visualize MA plot. Differentially accessible peaks for GATA1 high or low cells was used to calculate motif enrichment (getMotifEnrichment function in the ChrAccR R package) using the CIS-BP TF motif database (from chromVARmotifs package). Adjusted p value (q value) was converted to - log(q value) and top enriched motifs were plotted by -log(q value) and odds ratio.

### ChromVAR analysis

Raw insertions counts at relevant consensus peakset regions were RPKM normalized, log2 transformed, and quantile normalized. ChromVAR deviation scores were calculated on the log transformed count matrix using getChromVarDev function in the ChrAccR R package. Top variable TF motifs’ deviation scores were plotted using ComplexHeatmap R package.

### GATA1 footprinting analysis

To calculate GATA1 footprinting as a measure of GATA1 occupancy, we calculated Tn5 bias-corrected, normalized insertions centered at GATA1 motif sites across the GATA1 consensus peak set using the aggregateRegionCounts in the ChrAccR R package using the following parameters: countAggrFun = “mean”, norm = “tailMean”, normTailW = 0.1, kmerBiasAdj = TRUE, k = 6. To compare accessibility in K562 cells with high vs low GATA1 across GATA1 sites of different binding affinity, we identified the highest scoring GATA1 sequence motif within each consensus peak, binned all sites into 20 equal bins based on the GATA1 motif score, and calculated the GATA1 footprint. We then measured accessibility flanking the GATA1 motif as the area under the GATA1 footprint plot from −50bp to −10bp and from +10bp to +50bp. For each motif score bin, the fractional change in accessibility was calculated as the average difference in accessibility between GATA1-high and GATA1-low samples normalized to the accessibility in the GATA1-low samples.

### Mass cytometry experiment

BM enriched CD34+ cells were thawed and stained for viability using cisplatin protocol^33^. After quenching and washing in CSM (at 250G for 5min), cells were stained for surface markers before fixing in 1.6% paraformaldehyde and permeabilizing in 100% methanol for 10min at 4C. GATA1 and cleaved caspase antibodies were stained intracellularly before washing in CSM at 600G for 5min. Cells were then re-fixed in 1.6% paraformaldehyde and DNA intercalator^34^ before analyzing on Helios mass cytometer (Fluidigm). Supplementary Table 1 details antibodies used, their clones and the metal isotope channels they were conjugated to. Resulting FCS files from Helios run contains single cell protein level abundance for ∼1mil BM cells.

### Mass cytometry analysis

FCS files were gated on cytobank or cellengine platforms for cisplatin low (viability) and then gated for BM progenitors and conventional MEP as detailed above. Manually gated populations were exported into R and acsinh transformed with a cofactor of 5 and normalized from 0-1. CD34+/CD38+ gated BM progenitor mass cytometry data was then correlated across GATA1 and all assayed surface markers using Spearman correlation and plotted using corrplot R package with hierarchal clustering. Boxplots of GATA1 high and low BM progenitors were constructed using top 8% of GATA1-expressing cells and remaining cells respectively in order to match frequency of GATA1 positivity captured in sorting for InTAC experiment. Manually gated canonical MEP and deduced candidate populations were plotted for GATA1 abundance using ggplot2 boxplot function. The data-driven, backgating algorithm GateFinder^26^ was applied on all BM progenitors with candidate population as target with 2 gating step parameter and predicted gates were plotted as scatterplots using ggplot2.

### Colony-forming unit (CFU) assay

BM enriched CD34^+^ cells were stained with CD34-FITC, CD38-APC/Cy7, CD71-PE, CD33-PE/Cy7, and CD84-APC and sorted for our putative GATA-1 high erythroid progenitor subpopulation. We also stained and sorted cells for conventional MEPs (CD34-FITC, CD38-BV421, CD45RA-AF700, CD10-BV650, CD123-PECy7). Antibody panels in Supplemental Table 2. For viability, 7-amino actinomycin D (7-AAD) was used. Our putative population corresponding to GATA-1 high BM progenitors were gated as singlet, viable CD34^+^, CD38^+^, CD84^hi^, CD71^hi^, CD33^−^ cells. Cells were sorting using a BD FACS Aria II (BD Biosciences) and collected in IMDM 2% FBS for further CFU assays.

CFU assays were performed using the MethoCult™ H4435 Enriched (STEMCELL Technologies). Briefly, progenitor BM sorted cells were seeded (250 or 500 cells/well) into 6 well SmartDish™ (STEMCELL Technologies). After incubation for 14 days, at 37°C in 5% CO_2_, hematopoietic colony-forming unit were automated counted and analyzed by STEMvision™ Human (STEMCELL Technologies). Differentiation frequency was calculated for each sorted population by number of resulting colonies/numbers of starting cells seeded.

### scATAC processing and clustering

Raw data files were downloaded from Granja et. al.^4^ which had scATAC and scRNA seq carried out in parallel on PBMC, BM and CD34-enriched BM. Processing was done using the ArchR package^23^, where Harmony^35^ was used to batch correct and MAGIC^36^ was used to impute gene accessibility scores. Further processing including iterativeLSI and subsequent UMAP embedding was carried out using ArchR’s built-in functions of addIterativeLSI and addUMAP. Pre-determined population annotations (from scRNA seq) were integrated into the scATAC data using constrained integration of the scRNA seq data (addGeneIntegrationMatrix from ArchR which uses Seurat’s transferAnchor function). Populations were filtered to exclude more differentiated PBMC and BM populations such as B cells, T cells and monocytes and focus the analysis on BM progenitors relevant to erythropoiesis. Seurat’s FindClusters approach was used on dimensionality reduced iterativeLSI embedding to cluster the scATAC data and clusters were labelled using predicted populations from annotated scRNA seq integration. MACS2 was run on the different scATAC clusters and reproducible peakset was curated using the addReproduciblePeakSet function with (n+1)/2 reproducibility with a maximum of 500 peaks per cell.

Populations were compared across accessibility in consensus peaks using binomial test after binarizing data and correcting for TSS enrichment and log10(nFrags) bias in getMarkerFeatures function of ArchR. Features differentially enriched across populations were plotted in a heatmap with a FDR<0.1 and Log2FC>0.5 cutoff. Key motifs such as GATA1, CEBPA, GATA2, KLF1, KLF2, SPI1, RUNX1, IRF4 and IRF8 were plotted for chromVAR deviation scores as stacked histogram across populations using plotGroups function in ArchR.

### Bulk sample projection onto scATAC space

Bulk ATAC count matrix calculated from relevant consensus peakset regions was converted into a summarizedExperiment data class. This was projected into the scATAC UMAP space after calculating iterativeLSI on bulk samples simulated as single cells using the projectBulkATAC function from ArchR. Resulting simulated single cell ATAC UMAP projection from bulk data (250 cells simulated per bulk sample) was plotted along with the scATAC data in the original UMAP embedding using ggplot. Closest 500 scATAC cells to simulated GATA1 positive samples’ bulk projection was quantified using mahanoblis distance to combined bulk sample centroid in projected UMAP space.

### Erythroid trajectory analysis

Erythroid trajectory was quantified by fitting splines through the early progenitor, early erythroid, mid erythroid and late erythroid clusters using the addTrajectory function in ArchR and normalizing pseudotime between 0-100. Trajectory was plotted using the plotTrajectory function. The GATA1 positive sample demarcations on trajectory was found quantifying closest scATAC cells’ pseudotime values on trajectory.

Gene accessibility, expression and chromVAR motif deviation scores for cells across the trajectory were extracted using the getTrajectory function from ArchR and heatmaps of top variable features were plotted. Normalized line plots were constructed by extracting GATA1 positive scATAC counts across 100 pseudotime bins and plotting using ggplot. GATA1 positive scATAC cells (by protein expression) was defined as GATA1 positive restriction point on trajectory (65-75 on pseudotime scale) and subsequently before and after restriction point was binned (40-60 and 80-100 respectively) as per plotted gene expression inflection points.

### Differential scRNA analysis

Integrated scRNA seq data was filtered for the abovementioned 3 restriction point related bins (before, during and after) and imported into Seurat R package. Data was log normalized with a scaling factor of 10000 and scaled after. FindAllMarkers function was used to detect only enriched markers in the 3 bins with genes detected in at least 10% of the total number of cells at a foldchange threshold of 2 using ROC analysis. Top 20 markers for each bin was plotted as a scaled heatmap using DoHeatmap function.

### Graphic design

All figures were constructed in Affinity Designer and schematics constructed in BioRender.com.

### Data availability

All sequencing data is available at GEO accession GSE167934.

## Acknowledgements

We thank members of the Bendall and Greenleaf labs for their support and advice. This study was supported by grants from National Institutes of Health 1DP2OD022550-01, 1R01AG056287-01, 1R01AG057915-01, and 1U24CA224309-01 (to S.C.B), RM1-HG007735, UM1-HG009442, UM1-HG009436, 1UM1-HG009442, U2CCA233311, U54-GH010426, and U19-AI057266 (to WJG), and GM110050 and GM127295 (to ETK). This work is also supported by the Defense Advanced Research Project Agency (W911NF1920185 to W.J.G.) and a Stanford Cancer Institute-Goldman Sachs Foundation Cancer Research Award (to W.J.G). WJG is a Chan Zuckerberg investigator. AFC is supported by an NIH F32 postdoctoral fellowship (5F32GM135996-02). RB is supported by A*STAR National Science Scholarship (PHD) from A*STAR Graduate Academy, Singapore.

